# Mechanistic Insights into MLKL Activation via Allosteric Pathways Identified Through Molecular Models

**DOI:** 10.64898/2026.06.10.731390

**Authors:** Ricardo Ramirez, Viviana Monje

## Abstract

The mixed lineage kinase domain-like protein (MLKL) is responsible for plasma membrane (PM) permeabilization during the last stage of necroptosis, a pro-inflammatory programmed cell death pathway. MLKL contains three domains: the pseudokinase domain (PsKD), the brace region, and the four-helical bundle (4HB). Phosphorylation of PsKD at amino acids S345 and S347 is essential for MLKL function as it triggers conformational changes that expose the 4HB and enables protein oligomerization and subsequent PM permeabilization. Understanding the molecular mechanisms of this allosteric signal is critical for the rational design of ligands that modulate cell death via necroptosis as terapeutic alternatives for neurodegenerative and inflammatory disorder. We simulated the wild-type, phosphorylated, and mutant MLKL proteoforms to build a Markov state model that revealed three dominant macrostates corresponding to the open, transition, and closed conformations. Hydrogen-bond/Hydrophobic network analysis, with a novel clustering approach, identified a switch in the allosteric pathways that favors the open state. Based on this model, we identified and tested computationally an MLKL mutant that facilitates 4HB exposure, and could favor protein oligomerization and compromise plasma membrane integrity. We present a comprehensive all-atom molecular dynamics study to characterize the conformational changes that enable exposure of the 4HB at the molecular level.

---

Necroptosis is a form of programmed cell death pathway implicated in cancer, neurodegeneration, inflammation, and infectious diseases, making it a central focus of biomedical research over the past decade [1–3]. This pathway culminates with the disruption of the plasma membrane (PM) by the phosphorylated and oligomerized form of mixed lineage kinase domain-like protein (MLKL) [4–8]. Necroptosis results in PM rupture and the release of intracellular contents, triggering an inflammatory response. The molecular mechanism by which MLKL permeabilizes the PM and how to regulate its activity for therapeutic purposes remains to be elucidated. Molecular modeling approaches play key roles to bridge this gap in knowledge by providing atomistic level details of the forces that drive biological processes as well as facilitating the selection of experimental targets. We set to identify the allosteric pathways that govern the activation of MLKL by exposing its four-helical bundle (4HB) domain with a comprehensive study using molecular dynamics and probabilistic models.

MLKL activation, oligomerization, and translocation to the membrane are consistent during the last steps of necroptosis across species, albeit some differences due to amino acid sequence [9–11]. Blocking MLKL activation has been proposed as a potential treatment for neurodegenerative diseases, where necroptosis contributes to neuronal loss [12, 13], as well as for inflammatory diseases, where it promotes chronic inflammation [14–17]. Conversely, activating MLKL has been proposed as a cancer therapeutic, since many cancer cells evade apoptosis but can still be eliminated by necroptosis [18–20]. A fundamental understanding of the structural and mechanistic basis by which phosphorylation by RIPK3 activates the MLKL molecular switch is critically needed to inform the development of targeted therapies for diseases exacerbated by necroptosis-induced inflammation.

In this work, we use murine MLKL (PDBID: 4BTF) as a model to investigate the molecular basis of MLKL activation. Murine MLKL contains 464 amino acids organized into three domains: the four-helix bundle (4HB, amino acids 1–124), the brace region (125–189), and the pseudokinase domain (PsKD, amino acids 190–464). Phosphorylation of the PsKD by RIPK3 has been proposed to act as a molecular switch that exposes the 4HB and activates MLKL [1]. Although multiple phosphorylation sites have been reported [1, 3, 15], RIPK3 specifically phosphorylates S345 and S347 in the context of necroptosis. Mutational studies targeting these sites and additional PsKD amino acids demonstrate that perturbations destabilizing the PsKD can promote MLKL activation and oligomerization, even in the absence of RIPK3 [1, 3]. Together, these observations support a model in which PsKD destabilization allosterically exposes the 4HB; however, no high-resolution structure of MLKL in an activated, 4HB-exposed conformation exists, and the molecular mechanism underlying this allosteric transition remains unknown.

These findings motivate our focus on phosphorylation events at S345 and S347, as well as PsKD mutations [1, 3] hypothesized to bias MLKL toward an active, 4HB-exposed state . While mutational phenotypes suggest that PsKD destabilization is sufficient for activation, how such perturbations reshape the MLKL conformational ensemble and communicate with the 4HB at the molecular level has not been determined.

We present a comprehensive simulation study to characterize the conformational changes in MLKL due to phosphorylation, identify an allosteric pathway between the PsKD and 4HB, and determine the effect of PsKD mutations on 4HB exposure and subsequent MLKL activation. We leverage Markov state models (MSMs) to resolve conformational ensembles, steered MD and umbrella sampling to probe the energetics of 4HB exposure, and graph-theory analysis to probe allosteric pathways for MLKL activation. Our findings provide a framework for rational design of ligands that target these allosteric pathways, offering a strategy to select drug candidates that modulate necroptosis for therapeutic purposes.

## 1 Results and discussion

Current experimental studies on MLKL are mainly focused on mutations assays to determine its killing potential, but none examine the mechanical or molecular basis of the result. Here, we bridge these experimental observations with molecular simulations to elucidate how MLKL phosphorylation reshapes its conformational ensemble and promotes activation.

Mutational analyses have identified phosphorylation at S345 as a key determinant of MLKL-mediated necroptosis. Alanine substitution at S345 significantly reduces MLKL killing activity, whereas the double mutation S345A/S347A completely abolishes cell death; in contrast, mutation of S347 alone has little effect [3]. Additional perturbations, including phosphomimetic substitutions (S345D and S345D/S347D) and PsKD mutations such as Q343A and K219M, induce necroptosis even in the absence of RIPK3 [1, 3]. Structural analysis indicates that Q343A and K219M disrupt a stabilizing hydrogen bond between K219 and Q343 observed in the inactive crystal structure (PDBID: 4BTF), suggesting that PsKD destabilization is sufficient to bias MLKL toward an activated state.

Guided by these experimental observations, we performed all-atom molecular dynamics simulations of single MLKL proteoforms in aqueous solution to probe the conformational dynamics induced by phosphorylation and activating mutations. Specifically, we simulated phosphorylated MLKL at sites S345 (S345p), S347 (S347p), and S345/S347 (2pMLKL), as well as experimentally characterized activating mutations S345D, S345D/S347D (S345/347D), and Q343A, alongside the wild-type protein. Each system was simulated in triplicate, yielding a total of seven systems (Table S1), with at least 3 *µs* of aggregate sampling per system and a minimum of 1 *µs* per replicate, following the protocols described in the Methods section.

### 1.1 Markov State Model identifies three macrostates for 4HB exposure

To characterize the factors that contribute to MLKL activation, we built a Markov State Model (MSM) based on the following trajectories: (i) replica 0 of the 2pMLKL system, extended to 2.5 µs due to its spontaneous 4HB exposure event; (ii) replica 0 of wtMLKL; and (iii) replica 2 of wtMLKL, for a combined simulation time of 4.5 µs. These trajectories were selected to ensure balanced sampling of open and closed conformations—2pMLKL predominantly explores open states, whereas wtMLKL explores closed states.

Principal component analysis (PCA) of the concatenated trajectories revealed two major basins in the conformational space (Fig. 1A). The MSM combined with PCCA+ identified three metastable macrostates (Fig. 1B): cluster 0 (purple), cluster 1 (green), and cluster 2 (yellow). To structurally differentiate these macrostates, we defined the angle between the 4HB and PsKD domains using two vectors constructed from centers of mass (COM) of amino acids exhibiting low root-mean-square fluctuation (RMSF)(arrows are shown in Figure 1C as reference). Based on the mean interdomain angles the states correspond to the transition state (cluster 0, 70.5°), opened state (cluster 1, 92.3°), and closed state (cluster 2, 65.2°) states. This angle was used to characterize the three macrostates, but it was not used as an input feature for MSM construction or state assignment.

This structural characterization aligns well with the kinetic connectivity inferred from the MSM. Specifically, the propensity of the transition state to convert to the closed or open states, while direct transitions between the closed and open states are comparatively infrequent. Conformational exchange primarily occurs through the transition macrostate, which serves as the main pathway between the closed and open conformations.

### 1.2 MD trajectories sample the three-state MSM model

The MSM provides a robust framework to classify conformational states and quantify changes in populations across different MLKL systems. The structures of all simulation trajectories were projected onto the three-state MSM, as shown in Fig. 1D,E. wtMLKL predominantly occupies the closed state, with near-unity probability. Single-site phosphorylation at S345p or S347p decreases the occupancy of the closed state and increases the transition state population, while the open state remains minimally populated. Replica 0 of the 2pMLKL system exhibits substantial occupancy of both the transition and open states, whereas the remaining 2pMLKL replicas display reduced closed-state occupancy compared to the wtMLKL. For the mutant systems, S345D and S345/347D show a pronounced increase in the transition-state probability with little occupancy of the open state. The Q343A mutant remains predominantly in the closed state, although with slightly lower probability than wild type.

Across all systems, the MSM reproduces the relative probabilities of clusters observed in the simulations and consistently projects independent trajectories onto the same metastable states, indicating internal consistency. The open state is under-sampled in most cases due to finite simulation times and energy barriers in the conformational landscape rather than a deficiency of the MSM itself. The observed trends of decrease in closed-state occupancy and increase in transition-state occupancy for phosphorylated and mutant systems are consistent with expectations from experimental studies [1, 3].

**Figure 1:**
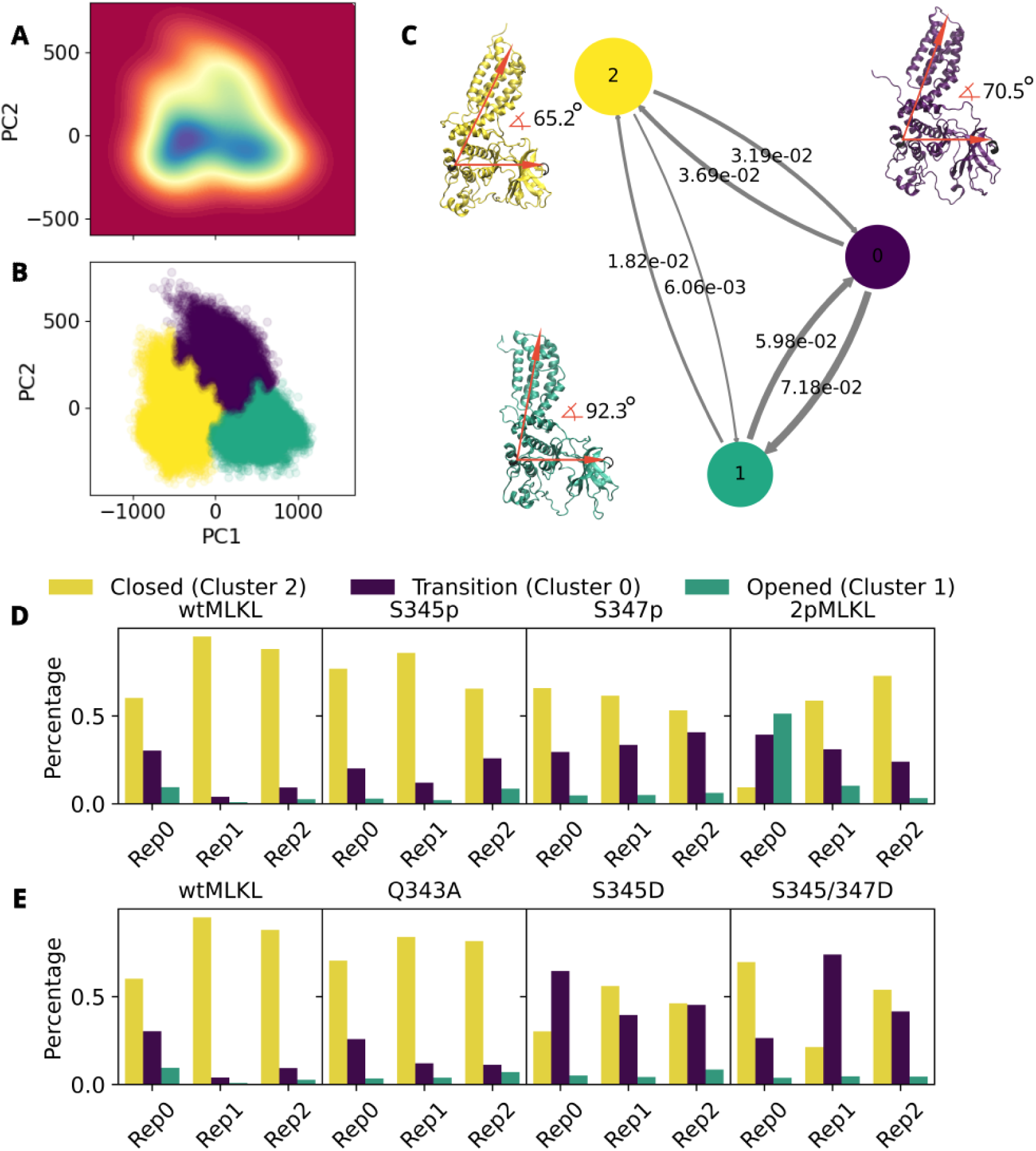
A. Density map of the two principal components of the concatenated trajectories B. PCCA assignment of frames to the transition (purple), open (green), and closed (yellow) MSM macrostates C. Transition rates for the three MSM macrostates D. Projection of the wtMLKL and the phosphoryled MLKL versions onto the MSM macrostates E. Projection of the wtMLKL and the MLKL mutants onto the MSM macrostates.

**Figure 2:**
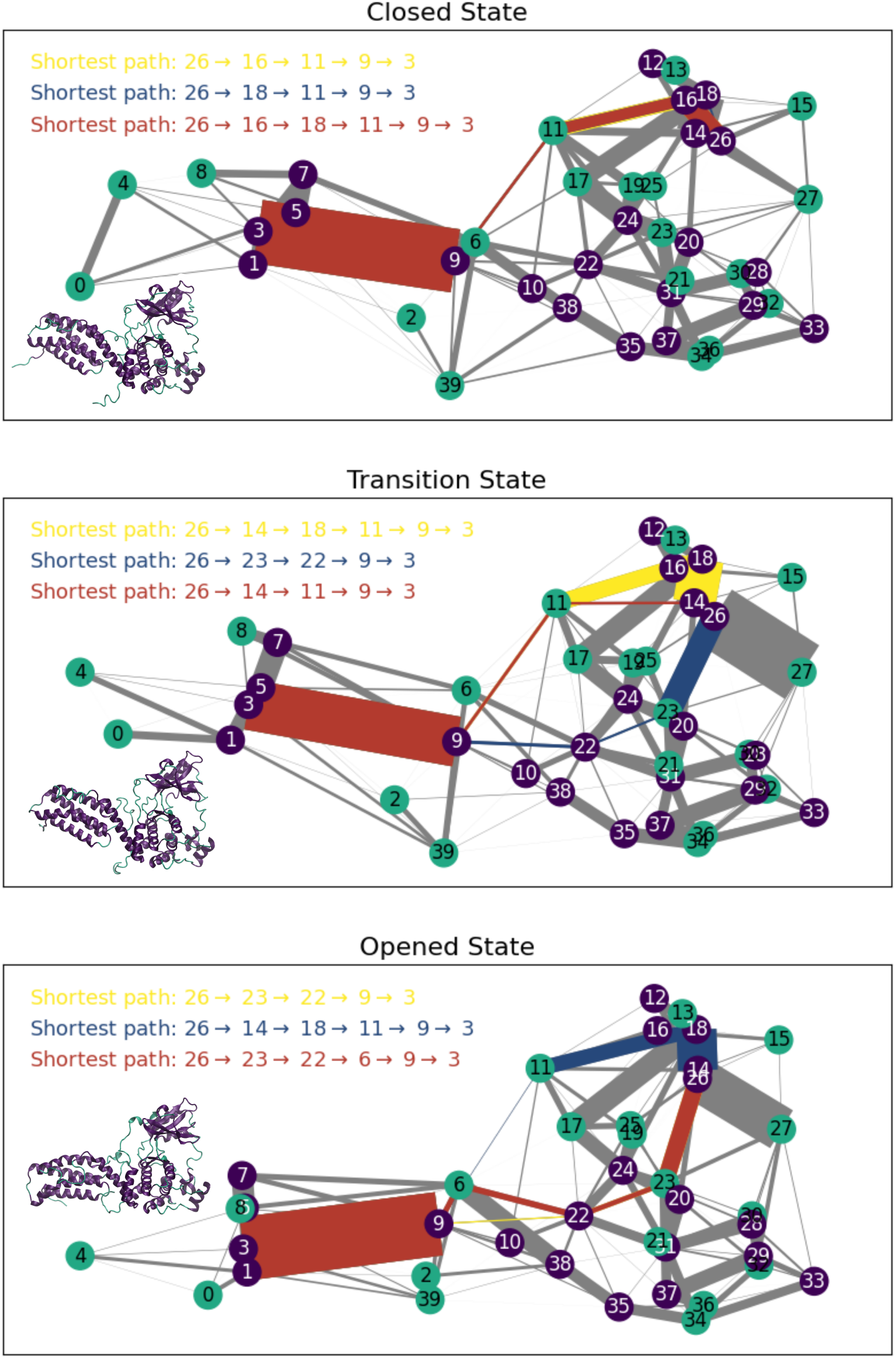
Coarse-grained graph network constructed with HB and HI interactions for the transition state (A), open state (B), and closed state (C). Nodes in green are amino acids in structured regions and nodes in purple are amino acids in disordered regions. The top three shortest paths in the network are listed in order of increasing length, yellow being the shortest.

### 1.3 Allosteric network modulates MLKL conformational switch

We analyzed the hydrogen-bond (HB) and hydrophobic interaction (HI) networks across the three MSM macrostates to determine the effect of phosphorylation on MLKL activation. Graph representations of HB networks have been used to reveal allosteric communication in large proteins, including the SARS-CoV-2 spike [21–23]. We extend this framework by incorporating weighted HI edges, whose interaction strength is approximately one-fifth that of HBs [24], to capture nonpolar contributions to stability. Each amino acid was treated as a node, with weighted edges representing the cumulative strength of HB and HI interactions (as shown in Figure S1). To enhance interpretability, a coarse-grained representation was developed to reduce the network from 469 to 40 nodes with edges corresponding to summed HB/HI interactions (Figure 2). Adjacent amino acids within the same secondary structure or a continuous disordered segment were grouped into single nodes to preserve the dominant communication routes while focusing on interactions between structural elements rather than individual amino acids.

Shortest-path analysis revealed distinct allosteric communication patterns across macrostates. In the closed state, the phosphorylation site (node 26) connects to the 4HB (node 3) through a route involving nodes 16, 11, and 9. In the transition and open states, alternative routes emerge through nodes 22 and 23, forming a new communication channel between the PsKD and the 4HB. These routes become increasingly dominant in the open state, suggesting that phosphorylation reorganizes intra-protein connectivity to facilitate 4HB release. Notably, node 23 corresponds to a disordered segment that forms a new contact with the phosphorylated helix of the PsKD (node 26), which is absent in the closed conformation. The emergence of this edge at nodes (23,26) likely mediates the conformational switch.

The networks display additional structural reorganization across macrostates besides the (23,26) connection (Figure 2). A new edge appears between nodes 2 and 38, specific to the transition and open states; this region overlaps with a helix implicated in MLKL inactivation as reported in recent experiments [25]. Interestingly, node 38 ranks consistently among the top-five central nodes in all graphs for both the open and closed states, based on closeness centrality (Table S2). Furthermore, node 38 connects directly to node 22, the most central node overall across all graphs. Shortest-path analysis between nodes 26 and 2 (Figure S2) shows that nodes 38 or 39 are consistently involved in both the open and closed states. Together, these results suggest that node 38 plays a key regulatory role in MLKL stability and activation, which support recent experimental findings [25].

Conversely, edges (6, 9), (9, 11) and (6, 17) markedly weakened in the transition and opened states, relative to the closed state. In Figure 3A–C) red edges mark some interactions that are strong in the closed state but weaken upon activation. Edge (6, 9) connects the disordered regions between helix 3 and 4 in the 4HB to the first helix in the brace. Edge (9,11) connects the first helix in the brace to a flexible region preceding the PsKD, whereas edge (6,17) links the disordered segment of the 4HB to one within the PsKD. These interactions appear to act as molecular tethers that stabilize the 4HB–PsKD interface; therefore, their loss during activation would facilitate 4HB disengagement. Four amino acids, S89, R137, E141, and D247, dominate these network changes (Figure 3D). To test the network-based prediction, we simulated an alanine mutant at these positions (S89A/R137A/E141A/D247A) for 600 ns and projected the trajectory onto the already trained MSM (Figure 3E). The mutant ensemble redistributed substantially across macrostates: the closed-state probability decreased closed to 0.2, while the transition state increased to 0.6. This shift, obtained without retraining the MSM, indicates that weakening these anchoring amino acids biases MLKL toward activation.

**Figure 3:**
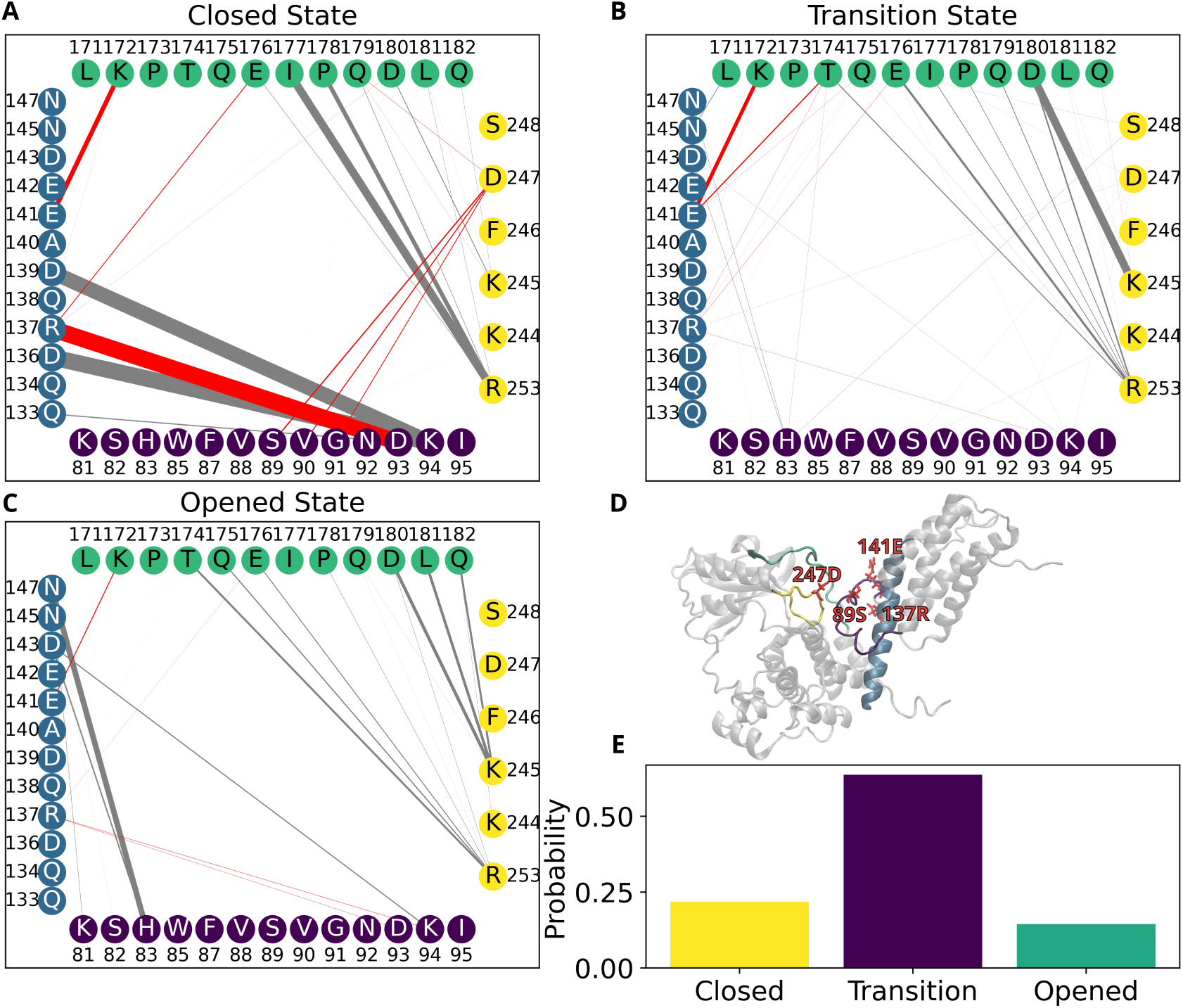
Dynamics of the edges anchoring the 4HB: (9,11) and (6,17) for the transition (A), open (B), and closed (C) states. Amino acids from nodes 6, 9, 11, and 17 are depicted in purple, blue, green, and yellow, respectively. (D) Structural view of the protein highlighting these nodes. (E) Projection of the mutated system onto the MSM.

This conclusion is consistent with the hypothesis that these edges constrain the 4HB in the inactive state, also in agreement with experiments [26, 27]. Mutation K172R [26], located at node 11, exhibits mild protection against necroptosis. Although a lysine-to-arginine change is conservative, arginine can form multiple hydrogen bonds that, in turn, enhance the (9,11) contact. According to our model, this subtle reinforcement would hinder 4HB release, aligning with the observed partial protection phenotype. Notice that although edges involving D139 are weakened in our model, we did not mutate this residue because previous studies show that a D139 V missense mutation already triggers necroptosis. This mutation is referred to as a “missense mutation”, however, based on our model, we predict that changes to D139 can potentially activate MLKL by weakening the edge (6,9)[27].

### 1.4 PsKD helices *α*2 and *α*5 are key for active states of MLKL

wtMLKL displays consistent dynamics across replicates, as indicated by the small error bars for the red curve on the RMSF profiles in Figure 4. The S347p system closely reproduces the wtMLKL profile, while 2pMLKL shows additional fluctuations at amino acids 339–347, which lies within node 25-26 in our coarse grain model. In contrast, the phosphorylated systems S345p and 2pMLKL exhibit pronounced increases in flexibility at amino acids 224–246, which are within nodes 14-16. Notably, these two segments include helices *α*2 (amino acids 228–243; node 16) and *α*5 (amino acids 339–347; node 26) in the PsKD, the same helices displaced by RIPK3 during MLKL activation, as reported by Xie et al. [28] and illustrated in Figure 4C,D. Hereafter, *α*2 and *α*5 are used interchangeably with nodes 16 and 26, respectively.

Our model identifies that the motion of the activation loop (*α*5, node 26) and the adjacent helix (*α*2, node 16) could promote the exposure of the 4HB. Replicate 0 of the 2pMLKL system shows that the activation loop *α*5 moves towards the solvent, resembling the RIPK3–MLKL co-crystal structure (PDBID: 4M69). This motion coincides with the strengthening of an edge between nodes 26 and 23 in the network model, reflecting that the interaction is facilitated by the loop displacement. These results are in agreement with studies by Garcia et al. [26], who reported that the activation loop becomes solvent-exposed upon phosphorylation by RIPK3.

It is important to note that all activating mutants (S345D, S345D/S347D, Q343A) display increased RMSF in *α*2 and *α*5 regions, whereas non-activating variants (347pMLKL, wtMLKL) remain stable. The limited sampling of the transition and open states in our MSM may stem from the slow timescale of helix rearrangement, which is difficult to capture in conventional all-atom MD. Nonetheless, the RMSF and network analyses consistently indicate that activation-prone systems share enhanced flexibility in *α*2 and *α*5, and that their concerted motion is a hallmark of MLKL activation.

### 1.5 Probing the energetics of helix-driven exposure of the 4HB

To test the hypothesis that the movement of helices *α*2 and *α*5 promotes protein activation, we performed steered molecular dynamics (SMD) simulations pulling the centers of mass of these helices toward their respective position in the RIPK3–MLKL co-crystal structure [28].

Figure 4E shows the projection of the SMD trajectories onto the three-state MSM, compared to unbiased simulations initiated from identical starting conformations. As expected, the closed state exhibits a reduced probability in all SMD trajectories, consistent with the hypothesis that coordinated movement of *α*2 and *α*5 facilitates 4HB exposure. The open conformation is favored in all systems except S345D/S347D, where the transition state is the most populated. Interestingly, for wtMLKL, the open state becomes dominant under steering conditions.

**Figure 4:**
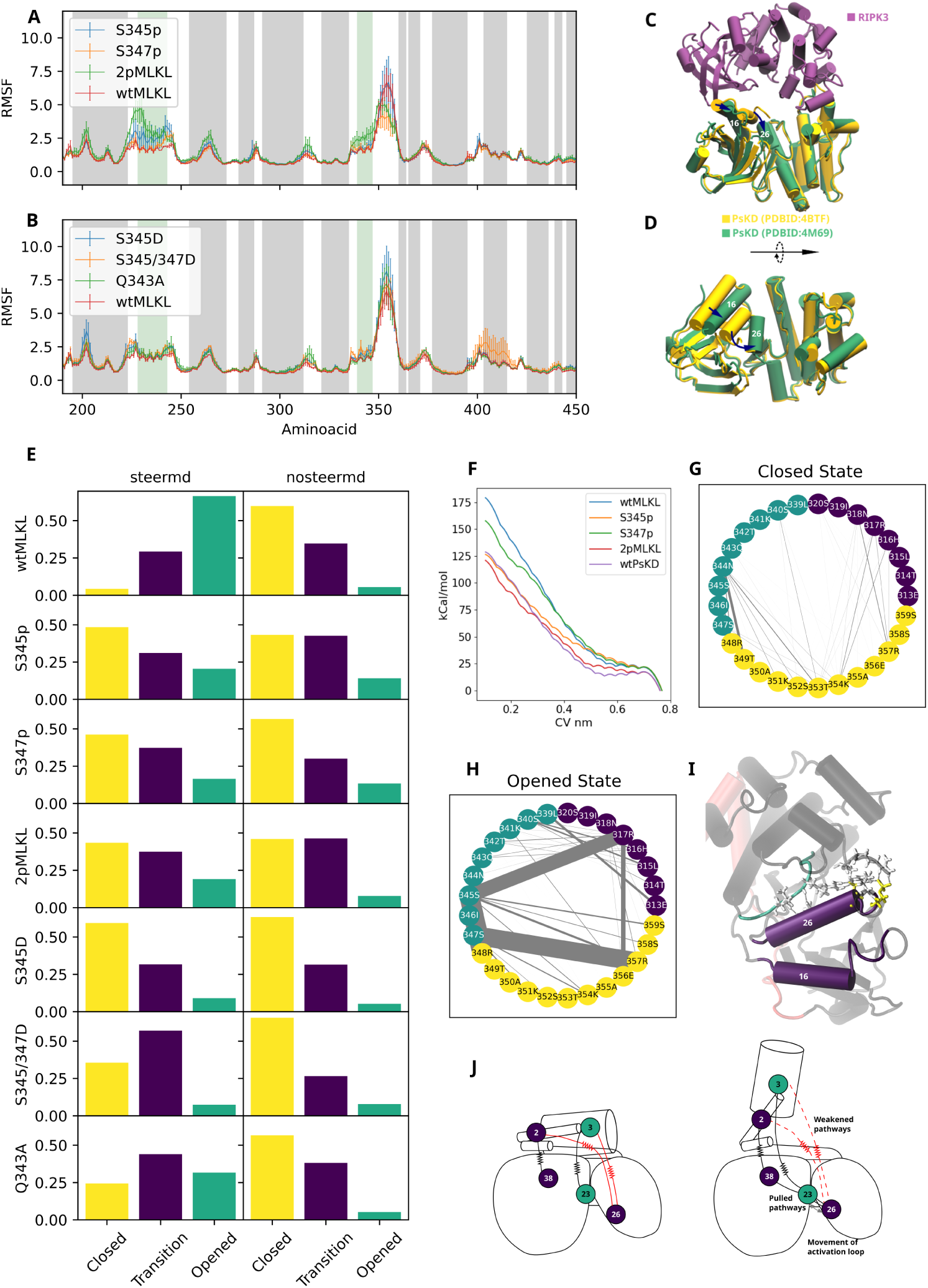
A. RMSF for the phosphorylated versions of MLKL B. RMSF for MLKL mutants C. Complex RIPK3(Purple)-PsKD(Green) from PDBID:4M69, compared to PsKD (yellow) from PDBID:4BTF D. Close-up of the PsKD in structures 4M69 and 4BTF E. Steer MD simulations projected to the MSM F. Umbrella sampling energy profiles for the movement of the helices *α*2 and *α*5 G,H. Isolated network for nodes 23, 26, and 27 for the closed and open states, respectively I. Close-up of the interactions of nodes 23,26, and 27 J. Proposed model for the switch mechanism.

This suggests that the displacement of *α*2 and *α*5, which RIPK3 directly contacts in the wild-type protein— may be sufficient to trigger 4HB exposure. In this view, phosphorylation of S345 and S347 would serve to stabilize the active conformation rather than initiate the structural rearrangement itself. In this view, phosphorylation is needed to keep the alpha helices displaced, or interacting with node 23.

### 1.6 The 4HB exposure switch arises from coupled helix motion and allosteric reorganization

Given that the movement of helices *α*2 and *α*5 facilitates 4HB exposure, umbrella sampling was used to compute the energetics of their displacement on several proteoforms. The resulting free-energy profiles reveal that wtMLKL exhibits the highest barrier to helix motion, followed by S345p, S347p, and finally 2pMLKL, which displays the lowest barrier (Figure 4F). Thus, phosphorylation progressively lowers the energetic cost of the conformational change, in agreement with experimental findings by Rodríguez et al., who reported that phosphorylation at both S345 and S347 sites combined produces stronger necroptotic activity than a single phosphorylation site at S345 alone.

To further examine 4HB exposure and the role of specific protein motions, we repeated the pulling simulations using a model containing only the wt PsKD (amino acids 190–464). Remarkably, the energy barrier for helix displacement in this truncated system decreased to the same level observed for 2pMLKL, indicating that this motion is an intrinsic property of the PsKD and forms part of the allosteric pathway leading to 4HB exposure.

Mechanistically, displacement of *α*2 and *α*5 enables the activation loop (node 26) to form new interactions with node 23, as revealed by the network model (Figure 3). In the open and transition states, the phosphorylated amino acids S345 and S347 display enhanced contacts with neighboring amino acids, including R317, R348, R357, S358, and S359 (compare Figure 4G with Figure 4H). Among these, the interaction between S345 and R317 is particularly strengthened, representing the primary connection with node 23. This interaction stabilizes the new conformation (Figure 4I) and facilitates activation. Mutating R317 to alanine would therefore be expected to reduce MLKL’s necroptotic potential, even upon phosphorylation.

We propose a mechanistic model for MLKL activation in which the closed state is characterized by a restrained 4HB connected to the PsKD by multiple allosteric “springs,” with the brace helix acting as a hinge. Upon phosphorylation or RIPK3 binding, the activation loop shifts toward node 23, weakening the dominant allosteric route that stabilizes the closed conformation (depicted as the red pathway in Figure 4J) and favoring new interactions between node 23 and nodes 1–7 (corresponding to the 4HB), thereby exposing the 4HB. The coordinated weakening and strengthening of these pathways exposes the 4HB through an allosteric molecular switch. Thus, our model provides a mechanistic basis for the proposed role of PsKD amino acids in triggering 4HB activation [1, 10, 28].

## 2 Conclusions

MLKL is the terminal effector of necroptosis and has been implicated in multiple pathological processes. Therefore, understanding its activation mechanism and identifying strategies to modulate its activity are fundamental for the rational design of new therapeutics. Through a combination of molecular dynamics (MD) simulations, Markov state models (MSMs), network analysis, and umbrella sampling, we established a comprehensive model that elucidates MLKL’s conformational switching mechanism, aligns with experimental findings, and predicts the functional consequences of specific mutations.

Our MD simulations explored the conformational landscape associated with the exposure of the four-helix bundle (4HB), revealing three dominant macrostates characterized using MSM. To interpret these states, we developed a novel coarse-grained network representation that groups continuous ordered and disordered regions into distinct blocks. These blocks interact through hydrogen-bond and hydrophobic contacts, maintaining the overall topology of the protein while drastically reducing network complexity. This approach enhances interpretability and allows direct mapping of network nodes to the amino acids driving key interactions. For instance, our model successfully explains experimental observations such as the D139 V mutation, previously classified as a missense mutation that constitutively activates MLKL. In our analysis, D139 establishes strong interactions with amino acids within the 4HB, and its mutation weakens these hydrogen bonds, destabilizing the anchoring of the 4HB and facilitating its exposure.

We propose this framework as a generalizable pipeline for studying proteins that undergo large conformational rearrangements. The workflow involves simulating conformational transitions via MD, clustering the resulting data using MSM, constructing hydrogen-bond and hydrophobic interaction networks, coarse-graining the networks through our block-based approach, identifying critical nodes and edges via centrality or shortest-path metrics, and mapping them back to specific amino acids.

Applying this methodology to MLKL revealed the molecular switch underlying its activation. The process begins with phosphorylation of S345 and S347 by RIPK3, which induces the displacement of two helices—particularly the activation loop. This loop establishes new contacts with two disordered regions: one adjacent and one internal. The interaction with the internal amino acids reorganizes allosteric communication pathways through the protein’s core while simultaneously weakening peripheral interactions responsible for tethering the 4HB. Consequently, this redistribution of network connectivity leads to the release of the 4HB, initiating membrane permeabilization.

In summary, our model provides a mechanistic description of MLKL activation that bridges structural dynamics with allosteric communication and highlights key amino acids whose perturbation may shift the protein toward active or inactive states. Beyond MLKL, this framework offers a powerful and generalizable strategy for dissecting activation mechanisms in other dynamic or allosteric proteins.

## 3 Methods

### 3.1 All-atom Molecular dynamics

We used all-atom molecular dynamics simulations to study three phosphorylated structures of MLKL based on PDBID 4BTF. Our systems, S345p, S347p, and 2pMLKL correspond to phosphorylated sites at S345, S347, and S345/S347. Additionally, we prepared mutated versions at S345D, S345D/S347D, and Q343A (hereafter referred to as S345D, S345/347D, and Q343A) for a total of seven systems, including wild MLKL as control. All systems were solvated with TIP3P water molecules and neutralized using K ions at 7.0 pH using the CHARMM-GUI solution builder [29–31], custom mutations and phosphorylations are done directly through this graphical interface. All systems were run in triplicates using the CHARMM36 m force field and periodical boundary conditions [32–34].

The equilibration runs were performed using GROMACS 2021.5 with an integration timestep of 2 fs [35]. Temperature was kept constant at 310.15 K using the Berendsen thermostat with a 1.0 ps coupling constant [36]. Pressure was set at 1 bar and isotropically controlled with the Berendsen barostat using a coupling time of 5.0 ps and compressibility of 4.5 *×* 10*^−^*^5^ [36]. All production trajectories were run using NPT dynamics on GROMACS, using the Nose-Hoover thermostat and Parrinello-Rahman barostat [37, 38] with the same coupling and compressibility settings as listed above. Non-bonded interactions were modeled using the Verlet force-switch function with cutoffs set at 1.0 and 1.2 nm for Lennard-Jones interactions [39]. Particle Mesh Ewald was used for long-range electrostatics and the LINCS algorithm to constrain bonds with hydrogen atoms [40]. Production runs were performed for 1us each, except 2pMLKL replicate 0 that was extended to 2.5us since it showed spontaneous 4HB exposure. All simulations in this article were run with resources available at the Center for Computational Research (CCR) at the University at Buffalo [41].

### 3.2 Markov State Model

To characterize the conformational landscape of MLKL and its modulation by phosphorylation, we constructed a Markov state model (MSM). Trajectories from the wtMLKL (replicas 0 and 2) and the phosphorylated MLKL (2pMLKL, replica 0) were concatenated, as these systems collectively span the conformational space associated with exposure of the 4HB during necroptosis. Principal component analysis (PCA) was then performed on two groups of structural features: (i) the pairwise distances between the *C_α_* atoms of the 4HB-PsK, 4HB-brace, and PsK-brace domains, and (ii) the PsK-4HB angles. The latter were computed in radians by defining a fixed reference point-the center of mass of amino acids 331-336-from which two sets of vectors were generated: those pointing toward the 4HB and those toward the PsK. The final angles were defined as the angles between these two vector groups. To minimize noise from flexible regions, only *C_α_* atoms belonging to secondary structural elements were included in the analysis.

The top two principal components were used for k-means clustering, yielding 80 microstates, which were then employed to build the MSM using Deeptime [42, 43] and validated using the Chapman Kolmogrov test (Figure S3). The Perron cluster analysis [44, 45] was subsequently applied to identify three kinetically distinct macrostates representing the main conformational ensembles of MLKL.

### 3.3 Network analysis

The initial network was build using a combination of hydrogen bonds (HB) and hydrophobic interactions (HI). HBs were computed using MDAnalysis with a 3.2 Å^2^ as a distance cutoff and 150*^◦^* angle cutoff. The results we comprised into a 469 *×* 469 matrix (accounting for the first five extra amino acids extra present in PDBID:4BTF), one entry per aminoacid where each value is the number of HB during the trajectory per total number of frames. As each pair or amino acids can have multiple hydrogen bonds, the entries can be greater than the unity. The HI were analyzed using MDAnalysis, counting an HI when the COM of two hydrophobic side chains is within 5 Å. The final result is a matrix 469 *×* 469 where each entry is the number of HI per total number of frames, which is then added to the HB matrix with a factor of one fifth to account for difference in strenght [24]. Networks were manipulated using networkx [46] from the adjacency matrices (HB+HI) obtained. In-house scripts were used for clustering continuous amino acids in ordered or disordered regions that comprised a single node. The shortest paths and node centrality were computed using the algorithm by Jin Y. Yen [47] and Freeman [48], respectively.

### 3.4 Steered molecular dynamics (SMD)

SMD simulations were performed starting from the equilibrated structure of the wtMLKL obtained from replica 0, with the goal of pushing two helices within the PsK domain to match their orientation in the crystal structure of the MLKL–RIPK3 complex (PDB ID: 4M69). Equilibrated MLKL coordinates were extracted from the 1 µs of unbiased production and solvated on CHARMM-GUI to ensure consistent starting configurations across all systems. The mutated and/or phosphorylated proteoforms were equilibrated following the procedures described in the AA-MD section.

A total of seven systems—including wild-type, phosphorylated, and mutant MLKL variants—were subjected to SMD simulations using PLUMED, chosen for its flexibility in defining collective variables (CVs) [49, 50]. The CV was defined as the distance between the COM of the targeted helices in the equilibrated MLKL structure and their corresponding helices in the crystal structure. This CV was driven using the MOVING_RESTRAINT module from 0.7 nm to 0 nm, with a linearly increasing force constant from 0 to 10,000 kJ mol*^−^*^1^ nm*^−^*^2^ over 4 ns, followed by a 100 ns phase with the constant force maintained.

To assess potential bias artifacts, all systems were also simulated for 100 ns under equivalent conditions but without the application of external forces, serving as control trajectories. The total simulation time for this data set is 1.4*µs*.

### 3.5 Umbrella sampling

To quantify the energetics associated with the movement of the PsK helices, umbrella sampling simulations [51] were performed using configurations obtained from the SMD trajectories described above. Thirteen sampling windows were generated along the collective variable (CV) in the interval of 0.7–0.1 nm. Initially, seven systems—corresponding to the wild-type, phosphorylated, and mutant MLKL variants—were simulated. In addition, we simulated only the PsK domain of the wild-type MLKL, resulting in a total of eight systems. Each window was simulated for 50 ns with a harmonic restraint of 12,000 kJ·mol*^−^*^1^·nm*^−^*^2^ applied to the CV. The first 10 ns of each trajectory were discarded for equilibration, and the potential of mean force (PMF) profiles were reconstructed using the Weighted Histogram Analysis Method (WHAM) [52].

