## Supplementary Information for "Mechanistic Insights into MLKL Activation via Allosteric Pathways Identified Through Molecular Models"

May 2026

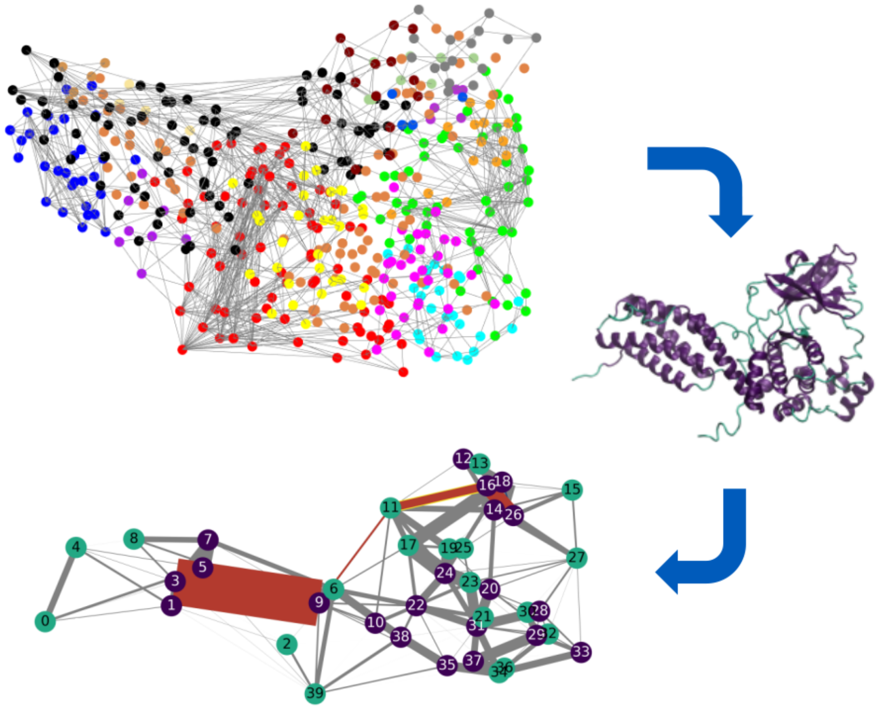


Figure S1: Illustration of the initial hydrogen bond plus hydrophobic interaction network and the rational for its coarse grain using the closed state.

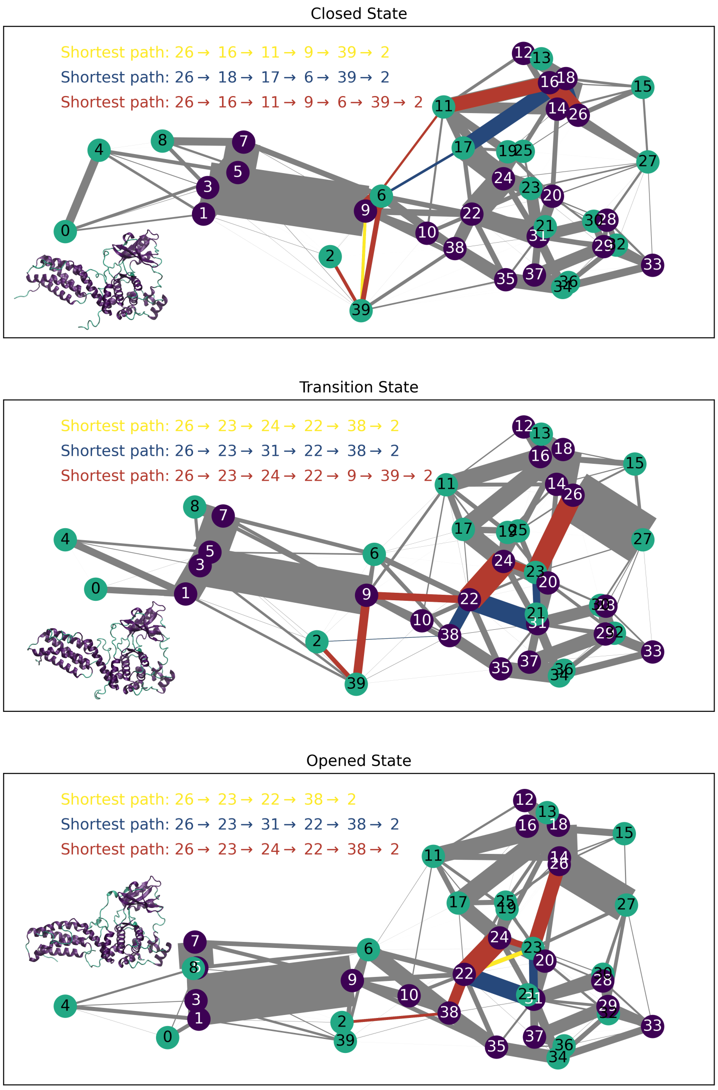


Figure S2: Network with shortest paths connecting the phosphorylation site and the disordered region of MLKL within the helices 1 and 2

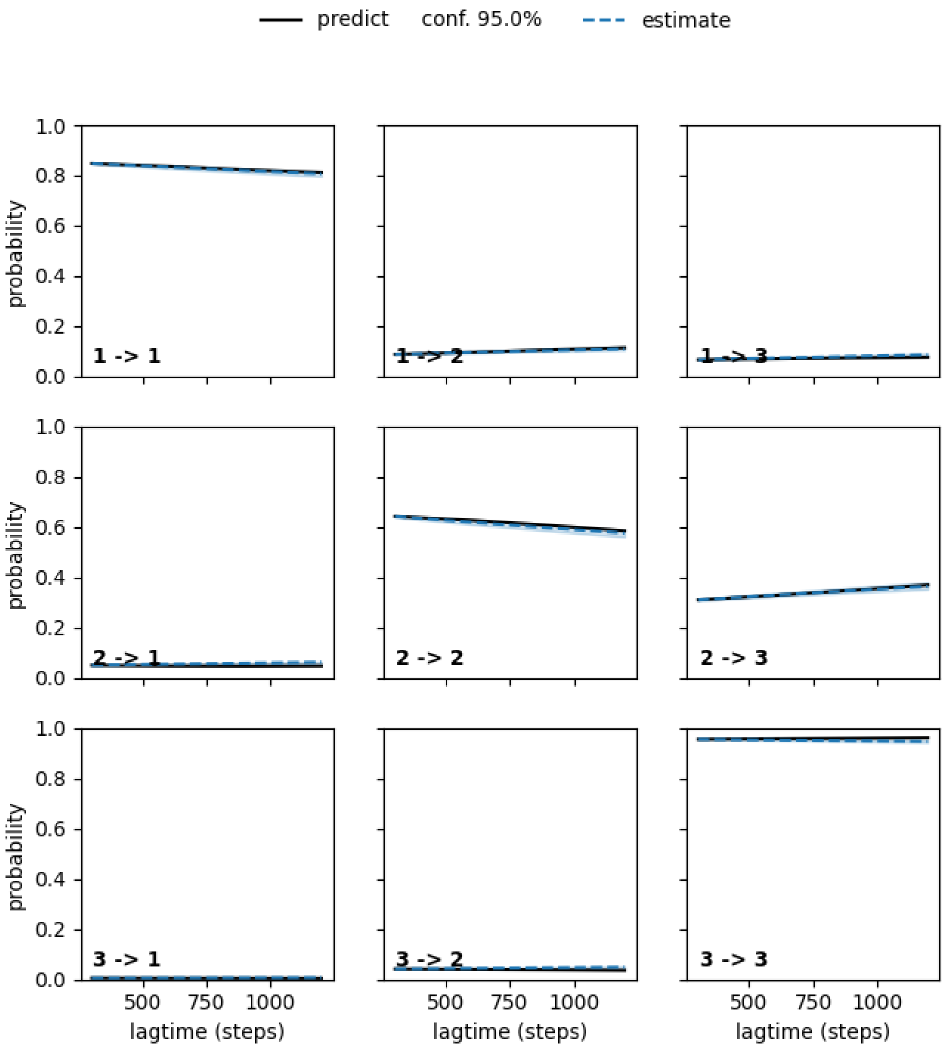


Figure S3: Chapman Kolmogorov Validation test for the Markov State Model

Table S1: Simulation details for all-atom MD systems, each run in triplicate for a total simulation time of 22.5 $\mu s$.

| Systems | Modification | Rep0 | Rep1 | Rep2 | Box size x,y,z |
|---|---|---|---|---|---|
| wtMLKL | – | 1$\mu s$ | 1$\mu s$ | 1$\mu s$ | 11.04, 11.04, 11.04 |
| S345p | Phosphorylation | 1$\mu s$ | 1$\mu s$ | 1$\mu s$ | 11.04, 11.04, 11.04 |
| S347p | Phosphorylation | 1$\mu s$ | 1$\mu s$ | 1$\mu s$ | 11.04, 11.04, 11.04 |
| 2pMLKL | Phosphorylation | 2.5$\mu s$ | 1$\mu s$ | 1$\mu s$ | 11.05, 11.05, 11.05 |
| S345D | Mutation | 1$\mu s$ | 1$\mu s$ | 1$\mu s$ | 11.05, 11.05, 11.05 |
| S345/347D | Mutation | 1$\mu s$ | 1$\mu s$ | 1$\mu s$ | 11.05, 11.05, 11.05 |
| Q343A | Mutation | 1$\mu s$ | 1$\mu s$ | 1$\mu s$ | 11.05, 11.05, 11.05 |

Table S2: Closeness centrality measures (showed in parentheses) for the coarse-grained nodes for the network presented

| Top centrality | Closed | Transition | Opened |
|---|---|---|---|
| 1 | 22 (0.52) | 22 (0.54) | 22 (0.52) |
| 2 | 24 (0.50) | 24 (0.52) | 24 (0.50) |
| 3 | 31 (0.47) | 31 (0.50) | 31 (0.49) |
| 4 | 17 (0.47) | 17 (0.47) | 38 (0.46) |
| 5 | 38 (0.46) | 9 (0.44) | 17 (0.45) |
| 6 | 23 (0.43) | 20 (0.44) | 20 (0.42) |
| 7 | 9 (0.43) | 18 (0.43) | 6 (0.42) |
| 8 | 20 (0.42) | 38 (0.43) | 18 (0.42) |
| 9 | 18 (0.42) | 3 (0.42) | 35 (0.41) |
| 10 | 19 (0.41) | 23 (0.41) | 34 (0.41) |
